# Reactivation-dependent transfer of fear memory between contexts requires M1 muscarinic receptor stimulation in dorsal hippocampus

**DOI:** 10.1101/2024.06.02.597068

**Authors:** Karim H. Abouelnaga, Andrew E. Huff, Kristen H. Jardine, Olivia S. O’Neill, Boyer D. Winters

## Abstract

**Background:** Memory updating is essential for integrating new information into existing representations. However, this process could become maladaptive in conditions like post-traumatic stress disorder (PTSD), when fear memories generalize to neutral contexts. Previously, we have shown that contextual fear memory malleability in rats requires activation of M1 muscarinic acetylcholine receptors in the dorsal hippocampus. Here, we investigated the involvement of this mechanism in transfer of contextual fear memories to other contexts using a novel fear memory updating paradigm.

**Methods:** Following brief re-exposure to a previously fear conditioned context, male rats (n=8-10/group) were placed into a neutral context to evaluate the transfer of fear memory. We also infused the selective M1 receptor antagonist pirenzepine into the dorsal hippocampus prior to memory reactivation to try to block this effect.

**Results:** Results support the hypothesis that fear memory can be updated with novel contextual information, but only if rats are re-exposed to the originally trained context relatively recently prior to the neutral context; evidence for transfer was not seen if the fear memory reactivation was omitted or if it occurred 6h prior to neutral context exposure. The transferred fear persisted for four weeks, and the effect was blocked by M1 antagonism.

**Conclusions:** These findings strongly suggest that fear transfer requires reactivation and destabilization of the original fear memory. Specific parameters likely dictate similar generalization in disorders like PTSD. The novel preclinical model introduced here, and its implication of muscarinic receptors in this process, should inform therapeutic strategies in this area.

## Introduction

Memory updating is a fundamental process that facilitates the integration of novel information into pre-existing memory traces, thereby maintaining their relevance and accuracy over time (Hupbach et al., 2007; Lee, 2009; McKenzie and Eichenbaum, 2011). This is crucial for adaptive behaviors, enabling individuals to adjust to changing environmental demands. However, this process could be exploited in the case of certain maladaptive memories, especially those rooted in fear or trauma. For example, a characteristic feature of contextual fear memories is generalization, whereby fear responses extend beyond the original fear-inducing stimulus to a broad spectrum of related stimuli, exacerbating the distress and dysfunction experienced by affected individuals (Dunsmoor et al., 2009; Lissek et al., 2014). The basis for such generalization remains unclear, but memory updating mechanisms could lead to a linkage of aversive memories to neutral contexts. It is therefore important to consider the environmental and neurobiological factors that could drive such associations.

Consolidated memories can become labile upon reactivation (Nader et al., 2000; Jardine et al., 2022), and this renewed malleability is related to underlying mechanisms of synaptic destabilization (Wideman et al., 2018; Lee et al., 2008). We have recently implicated the cholinergic system in reactivation-induced destabilization of object, spatial, and contextual fear memories in rats; specifically, M1 muscarinic acetylcholine receptor (mAChR) activity appears to be necessary for destabilization of these memory types (Abouelnaga et al., 2023; Stiver et al., 2015, 2017; Huff et al., 2022; Wideman et al., 2023). Furthermore, direct integrative updating of object memories with new contextual information was similarly blocked when M1 mAChRs were antagonized prior to reactivation of the original object memory (Jardine et al., 2020). These findings suggest that mAChRs are involved in the process of memory updating because of their permissive role in memory destabilization (Jardine et al., 2020). The goal of the present study was to demonstrate a similar behavioural process – and mechanistic basis – for the updating of contextual fear memories with novel, previously neutral, contextual information.

To this end, here we introduce a novel paradigm, the Fear Reactivation and Transfer (FRaT) task, which models fear memory updating and generalization in rats through the process of memory linking following the reactivation of a contextual fear memory. While there is evidence that the introduction of certain types of information, aversive or positive, can update fear memories (Grella et al., 2022; Haubrich et al., 2015, Zaki et al., 2023), to our knowledge, there is no task that models fear memory linking to other contexts, which is a hallmark characteristic of certain maladaptive memories. We hypothesized that the introduction of neutral contextual information immediately following fear memory reactivation leads to integration of such contextual information with the previously-consolidated memory, resulting in fear transfer. Moreover, given our previous report implicating dorsal hippocampal M1 mAChRs in contextual fear memory destabilization (Abouelnaga et al., 2023), we predicted that antagonizing these receptors should block the updating effect by preventing destabilization of the original contextual fear memory.

## Methods

### Subjects

75 male Long-Evans rats were obtained from Charles River, Quebec and arrived weighing between 150g and 250g. Testing began when rats reached an approximate weight of 275g. See supplementary information for additional details.

### Surgical Procedures

Rats in intra-cranial experiments were implanted bilaterally with 22-gauge indwelling guide cannulas (Plastics1; HRS Scientific, Quebec), targeting the dorsal hippocampus (dHPC). See supplementary information for details.

### Microinfusion Procedure

See supplementary information.

### Drug Administration: Intra-cranial

Pirenzepine (Sigma-Aldrich, Oakville, Ontario), a selective M1 mAChR antagonist was given in experiment 5. It was administered at a dose of 20 μg/μL 30 minutes prior to reactivation, which has previously been shown to block destabilization of object location memories (Huff et al., 2022). Physiological saline (0.9%) was used as a control (VEH).

### Fear Conditioning Apparatus

See supplementary information.

### Fear Reactivation and Transfer Task

The FRaT task (Fig. 1) consisted of three sessions run 24h apart. Prior to these, all rats were habituated to the alternate contexts (triangle and circle apparatuses) in counterbalanced order and on separate days for a period of 8 min in each apparatus. The triangle apparatus walls were made of white corrugated plastic, with a smooth black rubber floor. The posterior wall was 75 cm long, and the other two walls were 60 cm long; all walls were 60 cm tall. The circle apparatus was made of grey plastic, with a floor made of black fine-grain waterproof sandpaper, reaching 48 cm in height with a diameter of 53 cm.

**Fig. 1.**
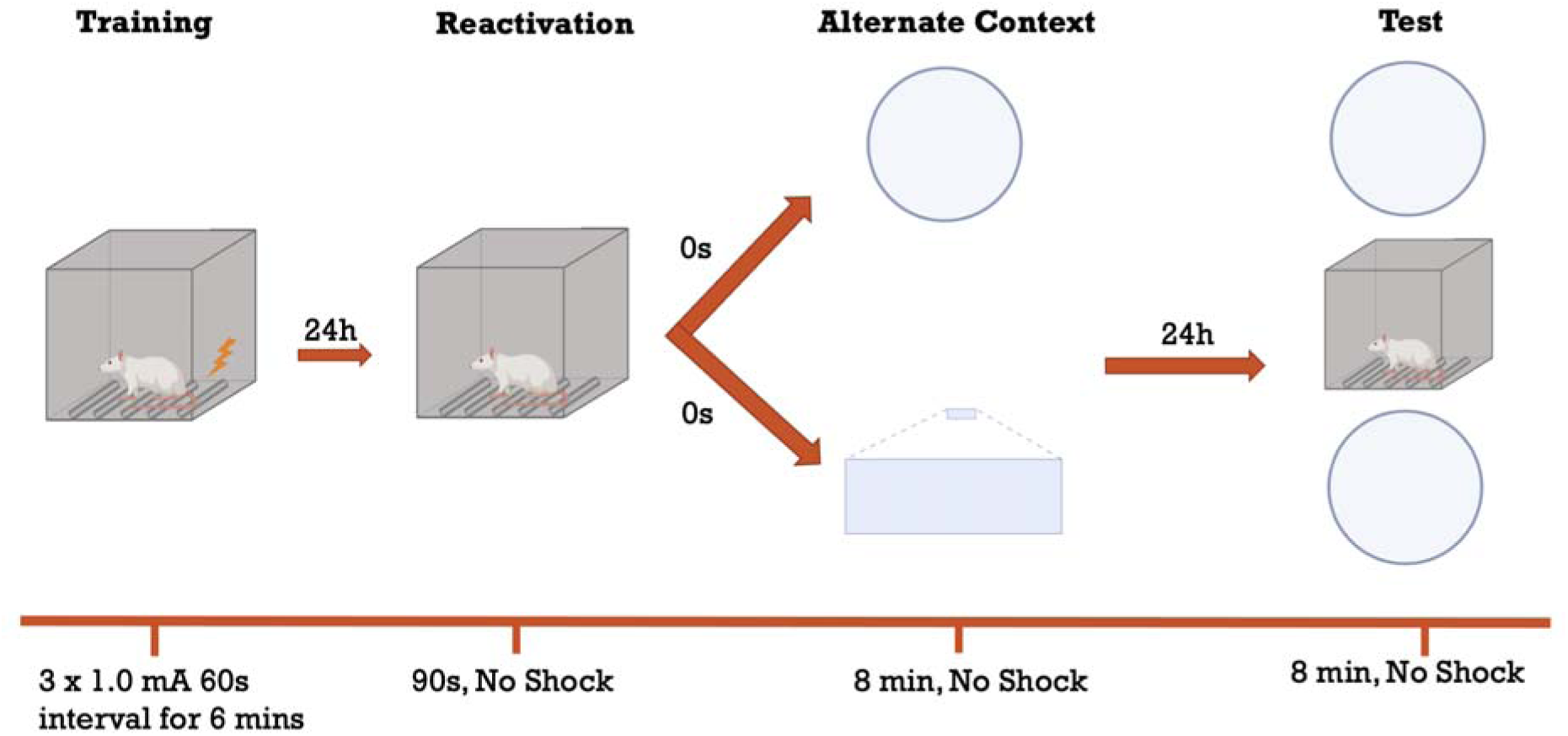
Illustration of the FRaT task. Rats undergo a training session in which they receive three foot-shocks. The reactivation session occurs 24h later and lasts 90s in the fear conditioning chambers. Animals are exposed immediately after to either a circular or triangular alternate context for 8 min before being returned to their home-cages. 24h later, they are assessed for freezing in either the same alternate context as the day before, a different alternate context, or the original fear conditioning chamber for a period of 8 minutes.

The training session took place 72 h following the second habituation day. For this session, the rats were placed in the fear conditioning chamber for a two min baseline period during which no foot-shocks occurred. Following this, three foot-shocks (1 mA, 1s each) were delivered 60s apart. This was followed by a 2-min post-shock period before the rat was removed from the box. The total training session time was 6 min.

Twenty-four hours later, the contextual fear memory was reactivated for each rat in its original training box by placement in the chamber for a 90-s period. There were no foot-shocks delivered in this session. Immediately after, the rats were placed in one of two alternate contexts, either the circle or triangle, counterbalanced, for an 8-min period during which they were free to explore; no foot-shocks were delivered.

The following day, the rats were tested for freezing in one of three contexts. They were either exposed to the same alternate context as the one they saw on the reactivation day, or a different alternate context. Alternatively, some rats were placed back into the fear conditioning box to assess the original memory.

## Experimental Approach

### Experiment 1

The purpose of this experiment was to determine whether a contextual fear memory can be updated with exposure to an alternate context following its reactivation. A group of 30 rats underwent a training session in the standard fear conditioning chamber before being divided into one of three groups based on the context in which they would be tested. The contexts were as follows (n = 10 each): same alternate context, different alternate context, or original fear chamber.

### Experiment 2

The purpose of this experiment was to determine whether the contextual fear memory updating that occurs in the FRaT task is reactivation dependent. For this experiment, 16 rats were used, and they were all tested in the same alternate context as the one they were exposed to post-reactivation. However, 8 of the 16 rats were not placed back in the fear chamber on Day 2; that is, half of the rats were exposed to an alternate context but did not first experience reactivation of the original memory.

### Experiment 3

The purpose of this experiment was to determine whether the updating contextual fear memory is persistent. For this experiment, the same cohort from Experiment 2 was retested 4 weeks later in the same alternate context.

### Experiment 4

The purpose of this experiment was to determine whether the contextual fear memory updating occurs within a restricted time period following reactivation. There is evidence that the reconsolidation window for fear memory updating lasts for less than 6h post-reactivation (Nader et al., 2000). For this experiment, the same FRaT protocol was used; however, groups of rats saw the alternate contexts either immediately after reactivation, 20 minutes after reactivation, or 6h after reactivation (n = 8 per group). All rats were tested in the same alternate context they experienced on the reactivation day.

### Experiment 5

The purpose of this experiment was to determine the necessity of M1 mAChRs by targeting them with intra-cranial infusions of the selective M1 mAChR antagonist pirenzepine into the CA1 region of the dHPC. 32 rats were obtained for this experiment. Rats were divided into one of four groups (Same-VEH, Different-VEH, Same-PIR, Different-PIR; Same = tested in same alternate context, different = tested in different alternate context). Each rat received a 2 min infusion of the drug or vehicle 30 minutes prior to the reactivation session.

### Data Analysis

Freezing behavior in the conditioning chambers was analyzed using Med Associates Inc’s stock software, Video Freeze. Freezing was defined as no movement except for that required for breathing (Maren, 2001). Freezing in alternate contexts was collected using a video camera situated above each of the alternate contexts. Freezing behaviour in the alternate context videos was analyzed using ezTrack (Pennington et al., 2019). See supplementary information for details.

## Results

### Histology

All rats that participated in behavioral testing in Experiment 5 were implanted with a bilateral cannula in which the tips were targeting the CA1 region in the dHPC. To verify placements, the rats were anesthetized with 0.8 mL of Euthansol (82mg/mL, Kirkland, Quebec) and then underwent pericardial perfusions with PBS and 4% formalin. Brains were then extracted and placed in a formalin solution for a minimum period of 24 hours prior to being transferred to a 20% sucrose solution for brain slicing preparation. Brains were sliced with a cryostat to 50 μm in width and then every third slice was mounted on a gelatin-coated slide. The slides were then thionin stained and visualized under the microscope. Placements were marked as individual dots, and a representative micrograph was obtained of the infusion tip in the CA1 region (Fig. 2).

**Fig. 2.**
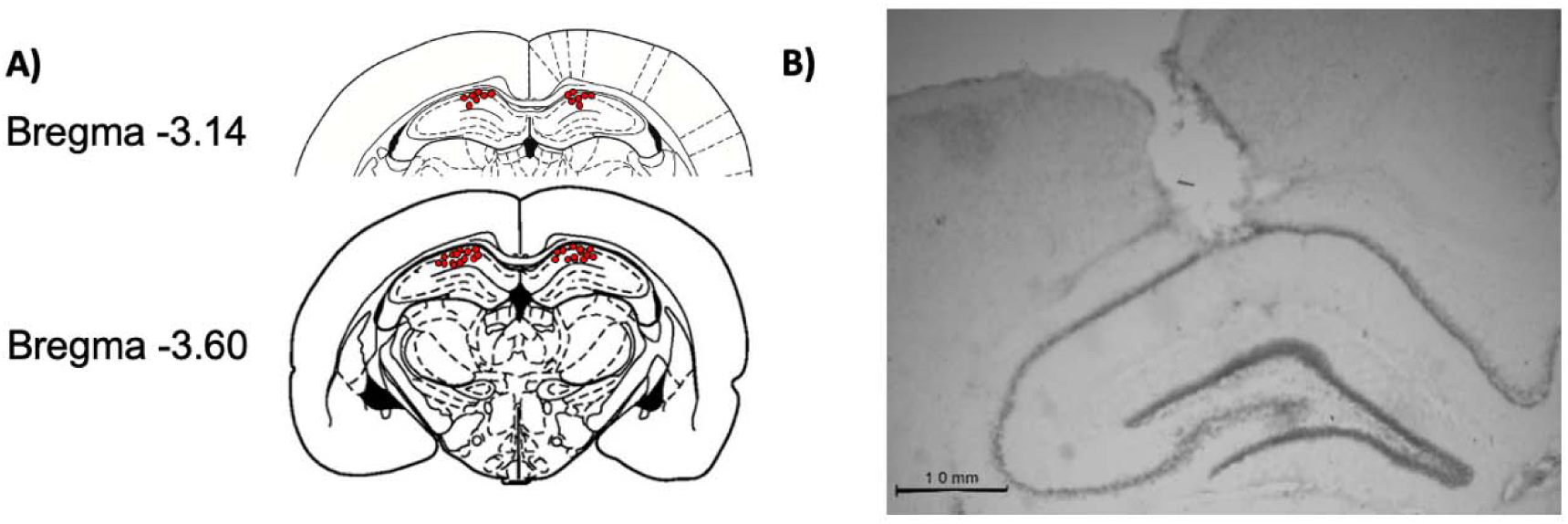
Schematic representation of the cannula placements for intra-dHPC infusions. A) Individual placements for each subject included in experiment 5 (n =21) B) Micrograph of an example cannula tip placement terminating in the CA1 region of the dHPC.

### Contextual fear transfers to a neutral context presented following reactivation of the original memory

The FRaT task was used in this experiment to determine whether contextual fear memories incorporate new contextual information if such information is presented immediately following reactivation. On reactivation day, results of a one-way between-subjects ANOVA showed no significant difference in freezing levels between the three groups (*F* (2,26) = 1.212, *p* = 0.313, η^2^ = 0.082; Fig. 3A). Additionally, there was no significant difference in freezing behaviour in the post-reactivation alternate context (*F* (2,26) = 1.391, *p* = 0.266, η^2^ = 0.093; Fig 3B)

**Fig. 3.**
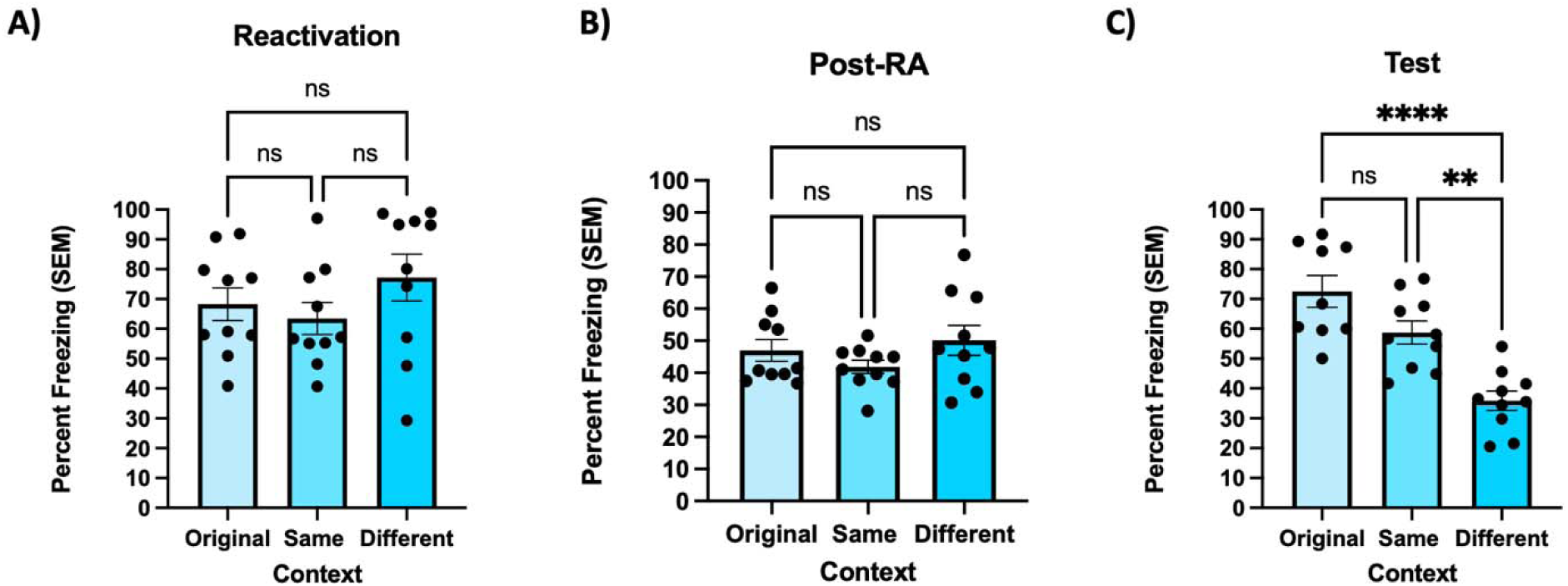
Contextual fear transfers to an alternate context when the original memory is destabilized. A) During the reactivation session, freezing levels did not differ between the three groups. Freezing behavior was measured as percent freezing ( SEM) across a 90-s reactivation session. B) There were also no differences in post-reactivation (Post-RA) context freezing. Freezing behaviour was measured as percent freezing ( SEM) across an 8-min post-reactivation session in the alternate contexts. C) In the test session, rats exposed to the same alternate context as the post-RA session (Same) froze significantly more than those tested in the different alternate context (Different). Same alternate context rats also froze similarly to the rats that were tested in the original fear conditioning chamber (Original). Freezing behavior (n = 8 per group) was measured as percent freezing ( SEM) across an 8-min test session. **** indicates p< 0.0001, ** indicates *p*<0.01.

During the test session, rats exposed to the same alternate context as in the post-reactivation session had significantly higher freezing levels compared to rats tested in a different alternate context (Fig. 3C). Results of a one-way between-subjects ANOVA indicated a significant effect of context (*F* (2,26) = 19.636, *p* <0.001, η^2^ = 0.602). Specifically, a Tukey’s multiple comparisons test revealed that rats exposed to the same alternate context on test day (Same) froze significantly higher than those tested in a different alternate context (Different) (*p* = 0.0015), but did not differ from rats placed back in the fear conditioning chamber where the original contextual fear memory was acquired (original) (*p* = 0.0699).

### Transfer of fear to a neutral context requires reactivation of the original fear memory

The purpose of this experiment was to determine whether reactivation of a contextual fear memory is necessary for transfer of fear to an alternate context. Here, only the ‘same’ condition was tested. One group underwent the reactivation session by being placed back into the fear conditioning chamber prior to being exposed to the alternate context (Same-RA), while the other group was placed into the alternate context without first being re-exposed to the training context (Same-NoRA).

Results of an independent-samples t-test showed no significant difference in freezing behavior between groups in the alternate contexts on reactivation day (*t*(14) = 1.455, *p* = 0.17; Fig. 4A). On test day, the group that received a reactivation session in the fear conditioning chamber froze significantly higher in the alternate context compared to the group that did not (*t*(14) = 3.348, *p* = 0.0048; Fig. 4B), suggesting that the fear transferred to an alternate context only when the original memory was reactivated.

**Fig. 4.**
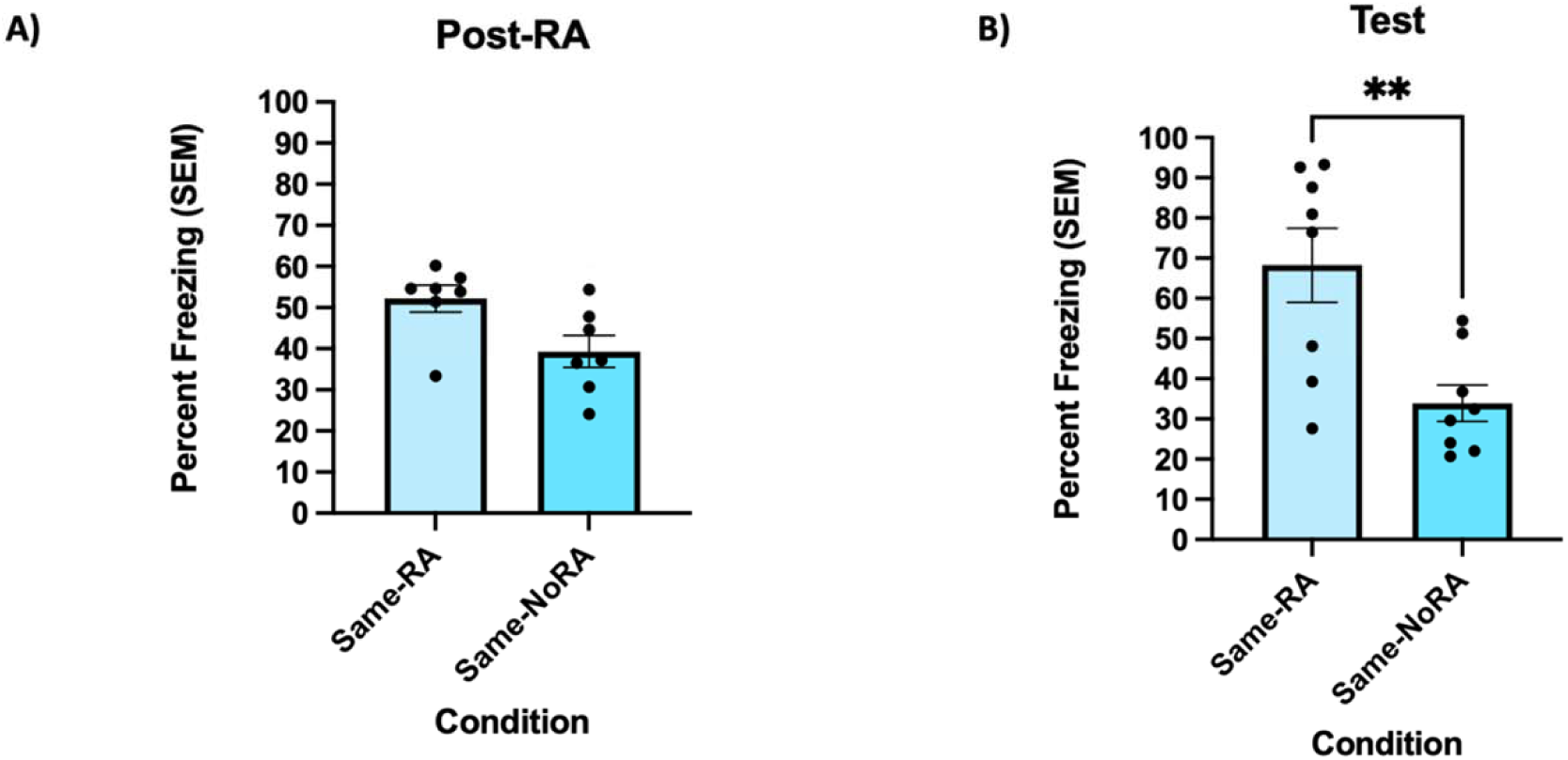
The transfer of fear to alternate contexts is reactivation-dependent. A) During the reactivation (RA) session, there was no significant difference in freezing behaviour between the groups in the alternate contexts. Freezing behaviour was measured as percent freezing ( SEM) across an 8-min post-reactivation session in the alternate contexts. B) For the test session, the group that was re-exposed to the fear conditioning chamber during the reactivation session (Same-RA) froze significantly higher in the alternate context than the group that was not exposed to the chamber (Same-NoRA). Freezing behavior (n = 8 per group) was measured as percent freezing ( SEM) across an 8-min test session. ** indicates *p*<0.01.

### Reactivation-mediated fear transfer persists for at least 4 weeks

To determine the persistence of the fear transfer effect in the FRaT task, the same rats underwent a retest session in the same alternate contexts that they were exposed to in the previous experiment. Results of an independent-samples t-test indicated a significant difference in freezing behavior between groups during the test session (*t*(14) = 3.202, *p* = 0.0064; Fig. 5), suggesting that the rats still experienced the alternate context as aversive even after 4 weeks.

**Fig. 5.**
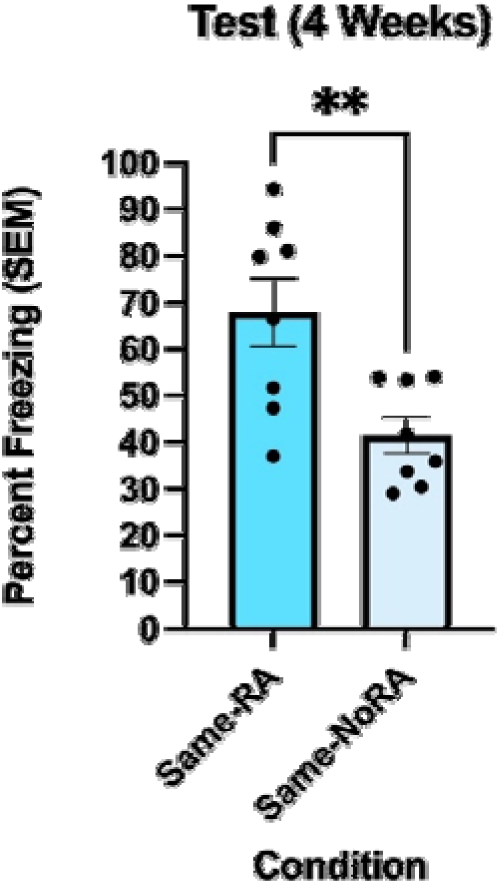
Transfer of fear to an alternate context is persistent when tested 4 weeks later. During the testing session, the Same-RA group from the previous experiment froze significantly higher than the Same-NoRA group, indicating that the transfer of fear to an alternate context is persistent. Freezing behavior (n = 8 per group) was measured as percent freezing ( SEM) across an 8-min test session. ** indicates *p*<0.01.

### Contextual fear memory updating occurs within a limited time window following reactivation

The purpose of this experiment was to determine whether this fear transfer effect only occurs within the putative reconsolidation window. Additionally, we aimed to determine whether this effect only happens if exposure to the alternate context occurs immediately after memory reactivation. Thus, we exposed rats to alternate contexts at one of three different delays: immediately after reactivation of the contextual fear memory (Immediate), 20 min after reactivation (20 min), or 6 h after reactivation (6h). On reactivation day, results of a one-way between-subjects ANOVA indicated no significant differences in freezing levels in the fear conditioning chamber (*F* (2,21) = 0.318, *p* = 0.731, η^2^ = 0.029; Fig. 6A). Additionally, there were no significant differences in freezing in the alternate contexts between the three groups (*F* (2,21) = 0.615, *p* = 0.550, η^2^ = 0.055; Fig. 6B).

**Fig. 6.**
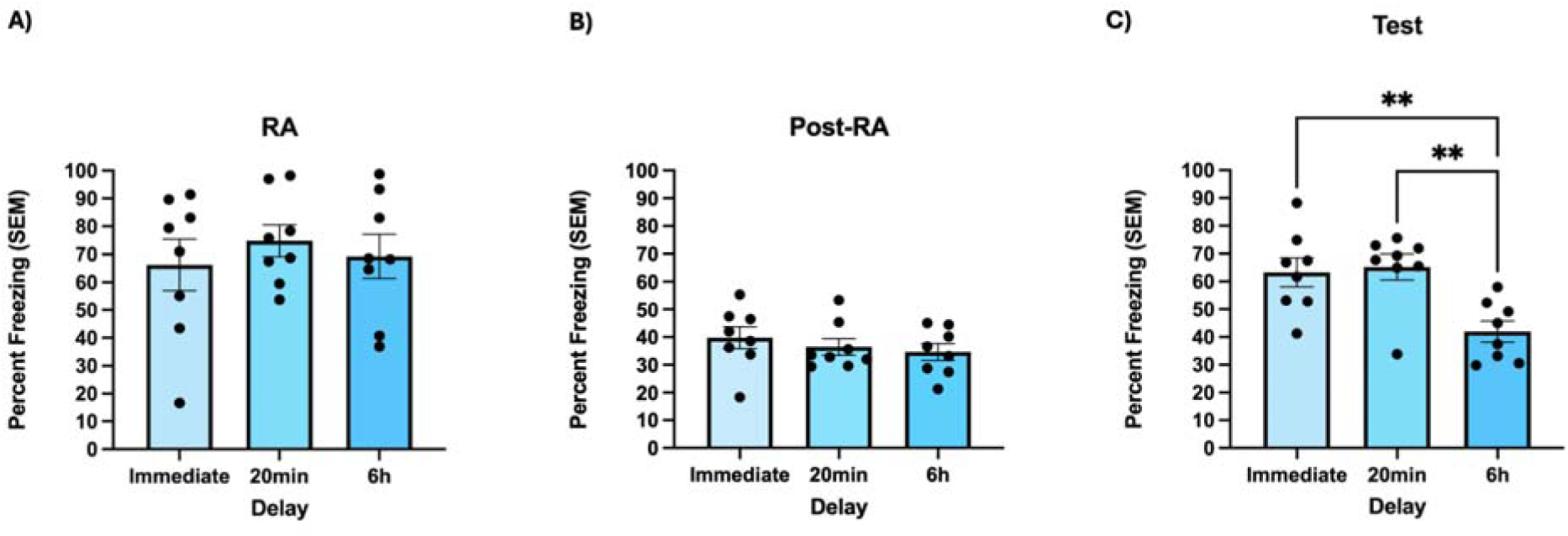
Fear memory updating only occurs if exposure to the alternate context happens within the 6h reconsolidation window. A) During the reactivation session, freezing levels did not differ between the three groups. Freezing behavior was measured as percent freezing ( SEM) across a 90-s reactivation session. B) There were no significant differences in freezing behavior during the post-reactivation sessions. Freezing behavior (n = 8 per group) was measured as percent freezing (± SEM) across the 8 min post-reactivation session. C) During the test session, the groups exposed to the alternate context immediately (immediate) or 20 min post-reactivation (20 min) froze significantly higher than the group exposed to the alternate context 6 hours (6h) post-reactivation. Freezing behavior (n = 8 per group) was measured as percent freezing (± SEM) across the 8-min test session. ** indicates p<0.01.

During the test session, rats exposed to the alternate context immediately or 20 min after reactivation froze significantly more than those that saw the alternate context 6h post-reactivation. A one-way between-subjects ANOVA revealed a significant effect of delay on freezing levels (*F* (2,21) = 7.987 *p* = 0.003, η^2^ = 0.432; Fig. 6C). Specifically, a Tukey’s multiple comparisons test revealed that rats exposed to the alternate context immediately post-reactivation (Immediate) froze significantly higher than those placed into the alternate context 6h later (6h) (*p* = 0.0091). However, there was no significant difference between the immediate group and the group seeing the alternate context 20 min later (20 min) (*p* = 0.9512).

### M1 mAChR antagonism in dHPC blocks reactivation-dependent contextual fear memory updating

Given the involvement of M1 mAChRs in the destabilization of contextual fear memories, as well as the role of the dHPC in this process (Abouelnaga et al., 2023), we predicted that the updating of contextual fear memories would also be dependent on dHPC M1 mAChR stimulation. Here, we directly antagonized these receptors in the dorsal CA1 during memory reactivation. On reactivation day, there were no significant differences between all groups (F (3,20) = 0.255, p = 0.857, η2 = 0.037). Additionally, there was no significant effect of pirenzepine on reactivation day freezing levels in the fear conditioning chamber (F (1,20) = 0.479, p = 0.497, η2 = 0.023; Fig. 7A). Post-reactivation freezing in the alternate context was not significantly affected when pirenzepine was infused 30 minutes prior to reactivation (F (1,20) = 0.617, p = 0.442, η2 = 0.030), nor was there an effect of post-reactivation context (F (1,20) = 0.208, p = 0.653, η2 = 0.010; Fig. 7B). Additionally, there was no significant interaction between the drug infused pre-reactivation and the context the rats were exposed to post-reactivation (F (1,20) = 0.298, p = 0.591, η2 = 0.015).

**Fig. 7.**
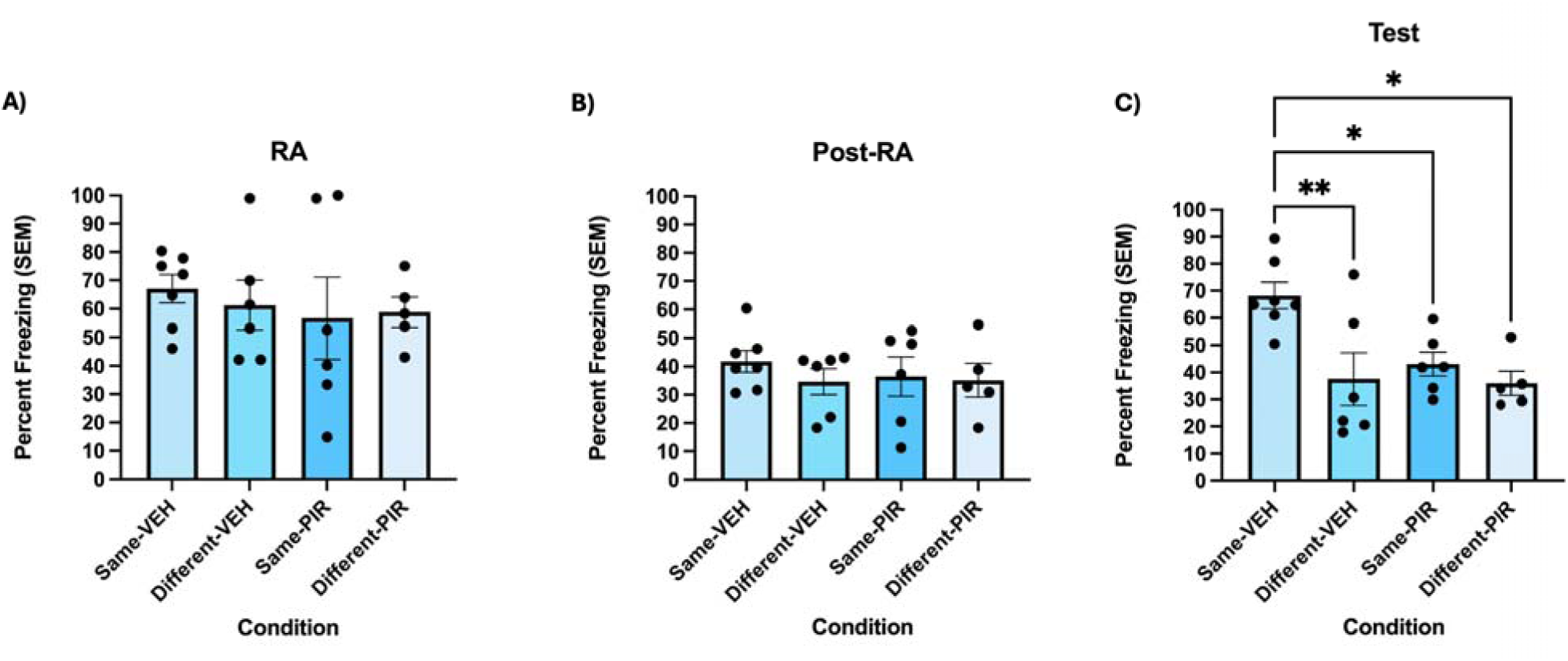
Contextual fear memory updating requires M1 mAChR activity in the CA1 region of the dHPC. A) During the reactivation session, freezing levels did not differ between the three groups. Freezing behavior was measured as percent freezing ( SEM) across a 90-s reactivation session. B) There was no significant difference in freezing behaviour during the post-reactivation session. Freezing behavior was measured as percent freezing (± SEM) across the 8 min post-reactivation session. C) The Same-PIR group froze significantly less than the Same-VEH group during the test session, implying that PIR disrupted the updating of the fear memory to incorporate alternate contextual information. Freezing behavior was measured as percent freezing (± SEM) across the 8-min test session. * indicates p<0.05, ** indicates p<0.01.

During the test session, a 2×2 between-subjects ANOVA revealed a significant interaction between drug infused prior to reactivation and the context that the rats were exposed to post-reactivation (F (1,20) = 3.349, p = 0.004, η2 = 0.476; Fig. 7C). Post-hoc comparisons indicated that the Same-PIR group froze significantly less than the Same-VEH group (p = 0.0393), implying that PIR blocked fear memory updating in the same alternate context condition.

## Discussion

In this investigation, we introduce a novel approach to studying the transfer of learned fear to other contexts, with a specific emphasis on the importance of reactivation of the original memory and the role of M1 mAChRs. Our findings, derived from the Fear Reactivation and Transfer (FRaT) task, suggest that novel contextual information can be incorporated into an existing fear memory and that this memory modification is reactivation- and time-dependent.

First, we used the FRaT task to update contextual fear memory in rats, incorporating information from a neutral context presented shortly after reactivation of the original fear memory. This finding is consistent with reconsolidation theory, which posits that upon reactivation, memories temporarily enter a labile state, making them susceptible to modification (Nader et al., 2000). We have previously described a similar phenomenon for object memory updating in rats (Jardine et al., 2020). Updating of contextual fear memory with new information has been explored in other studies investigating the linking of aversive experiences across time (Jung et al., 2023; Zaki et al., 2023). Furthermore, others have demonstrated the possibility of updating an established contextual fear memory with appetitive information, suggesting the versatility and adaptability of memory processes (Ramirez et al., 2013; Haubrich et al., 2015). However, here we demonstrate updating of a contextual fear memory without altering the original conditioned response, as seen in Experiment 1 where rats were still freezing in the fear conditioning chamber on test day even if alternate contextual information had been introduced post-reactivation. Moreover, the present findings appear to be the first to demonstrate the spread of fear from one context to another according to reconsolidation-based mechanisms.

Behaviour consistent with contextual fear transfer was only seen in the FRaT task when rats were exposed to the original conditioned context shortly prior to alternate context presentation; no updating occurred when the original context exposure was omitted or when the reactivation occurred 6h prior to alternate context exposure. The reactivation- and time-dependence of this effect is significant as it delineates the specificity of the memory updating process. It suggests that the rats are not indiscriminately generalizing their fear to any context they encounter, but rather, the reactivation primes the original memory for subsequent modification. This distinction between memory updating and generalized fear is crucial for understanding the nuances of memory processes. If the rats were simply generalizing their fear to any context through fear memory updating, it would imply a broad and non-specific response. However, our results indicate a more targeted and specific process, whereby the reactivated original memory is modified to incorporate new information presented within a restricted period of lability (Nader et al., 2000). It is also worth noting that rats in the various conditions did not differ in terms of freezing either during the memory reactivation session or when they were placed into the alternate contexts following reactivation; this pattern reinforces our interpretation of the present results in terms of a specific memory reconsolidation-based process of fear transfer.

Additionally, our results showing that freezing in the alternate context persists when tested 4 weeks later suggest a lasting, rather than temporary, effect. A shorter-term effect might indicate a less specific generalization related to the recent exposure to the original context. This alternative explanation is also not supported by our failure to see enhanced freezing when rats were tested in the different alternate context condition. Finally, the fact that fear spreads to the alternate context when it is presented 20 min following exposure to the original context, while consistent with reconsolidation theory, suggests that the effect observed in the FRaT task is not the result of backward second-order conditioning, which would likely require a much shorter delay between context presentations (Barnet et al., 1991; Mowrer et al., 1988).

The neural mechanisms underlying the present effect remain unclear. However, linking of memories through co-activation of neuronal ensembles seems a likely explanation. Cai et al. (2016), for example, demonstrated that memories for contexts encoded closely in time (5h) could be linked apparently via overlapping neural ensembles. Furthermore, Zaki et al. (2023) report that strong fear conditioning in one context can lead to later offline reactivation of the neural ensemble associated with the trained context as well as that for a previously presented (2 days earlier) neutral context, resulting in retrospective linking of the learned fear with that neutral context. While a similar process might be at play in the current study, the present behavioural effects are unique in that they demonstrate a critically reactivation-dependent form of prospective memory linking. Indeed, fear transfer in the FRaT task was not observed if the original contextual fear memory was not explicitly reactivated. The explicit reactivation session used here could accomplish something similar to the ensemble co-activation implicated by Zaki et al (2023), but possibly in a more selective and prospective manner. The effect reported by Zaki et al (2023) was only observed retrospectively and not seen when a relatively low shock protocol was used for the initial fear conditioning. Here, fear transfer occurred in a highly reactivation- and time-dependent manner. Moreover, despite both alternate contexts being pre-exposed 72h prior to the fear conditioning session (to habituate rats to these contexts), fear only transferred to the specific alternate context presented following re-exposure to the trained context. Thus, a generalized linking of fear with contexts explored within a few days of the training event did not occur; the original fear memory was only updated with the specific contextual information presented within a specific window of time following memory reactivation. There is one earlier report of apparent memory linking following reactivation, whereby fear conditioning to a second tone was enhanced when conducted shortly after presentation of a different tone previously paired with shock (Rashid et al. 2016). In contrast, however, here we provide evidence for memory reactivation- and time-dependent transfer of a fear response to an otherwise neutral context.

The fear memory updating effect reported here requires mAChR stimulation. Here, we targeted the dorsal CA1 with a selective M1 mAChR antagonist to block destabilization of the contextual fear memory when reactivated in the original fear conditioning chamber. The fact that pirenzepine blocked reactivation-dependent transfer of fear to the alternate context is consistent with the implication of M1 mAChRs in many forms of memory destabilization (Stiver et al., 2015,2017; Huff et al., 2022; Wideman et al., 2023; Abouelnaga et al., 2023), as well as our current hypothesis that a reconsolidation-based mechanism underlies the behavioural effect in the FRaT task. Our past work suggests that M1 mAChR activity during memory reactivation could lead to downstream activation of CaMKII and the ubiquitin proteasome system (Stiver et al., 2017; Wideman et al., 2023), consistent with a documented role for synaptic protein degradation in memory destabilization (Lee et al., 2008; Jarome et al., 2011). The plasticity resulting from such a process, combined with co-activation of contextual representations (ensembles) in the brain could facilitate updating of the original fear memory with the new contextual information, leading to fear transfer. Thus, M1 mAChRs could represent a therapeutic target for the prevention of fear generalization in disorders such as PTSD.

The FRaT task represents a novel animal model to facilitate the study of fear memory generalization and its behavioural and biological bases. To this end, the current study presents clear avenues for further research into the role of the cholinergic system, specifically M1 mAChRs, in fear memory updating. Elucidating memory modification mechanisms in this context has potential implications for better understanding and treatment of conditions like PTSD in which maladaptive memory generalization is very common and severely debilitating. The present results suggest that such generalization could occur through processes involved in reactivation and updating of the original traumatic memory and that interventions targeting this process could be effective in preventing the unfortunate spread of fear to stimuli not originally associated with the initial episode.

## Funding

This research was supported by a Natural Sciences and Engineering Research Council of Canada (NSERC) Discovery Grant (400176) held by BDW, Ontario Graduate Scholarship held by KHA, NSERC Canadian Graduate Scholarship – Doctoral (CGS-D) held by AEH, NSERC Canadian Graduate Scholarship – Doctoral (CGS-D) held by KHJ, and Ontario Graduate Scholarship held by OSO

## Authorship Contribution Statement

Karim H. Abouelnaga: Conceptualization, Methodology, Investigation, Formal analysis, Writing - original draft, Writing - review & editing. Andrew E. Huff: Investigation, Formal Analysis. Kristen H. Jardine: Conceptualization, Investigation. Olivia. S. O’Neill: Investigation. Boyer D. Winters: Conceptualization, Methodology, Writing – review & editing, Supervision, Funding acquisition.

## Disclosures

The authors report no financial interests or potential conflict of interests. The current manuscript will be submitted to the preprint server bioRxiv.

## Notes

### Competing Interest Statement

The authors have declared no competing interest.

