## Supplemental Methods for "Reactivation-dependent transfer of fear memory between contexts requires M1 muscarinic receptor stimulation in dorsal hippocampus"

### **Subjects**

75 male Long-Evans rats were obtained from Charles River, QC and arrived weighing between 150g and 250g. Testing began when rats reached an approximate weight of 275g. Upon arrival, they were placed in standard rectangular cages (48 x 26 x 20cm) composed of polycarbonate and containing Envirodry bedding and standard enrichment (nesting materials and wooden block). They were housed with a reversed light/dark cycle in which (lights off 8:00-20:00). Rats were acclimated to the room for one week before experiencing researcher handling in preparation for the experiments. Food and water were available *ad libitum*. Behavioral testing took place during the dark phase. All procedures were approved by the Animal Care Committee of the University of Guelph.

**Surgical Procedures**

For experiment 5, each rat was implanted with cannulas in the dHPC. Twenty-two gauge indwelling guide cannulas (Plastics 1, HRS Scientific, Quebec) were implanted bilaterally in the CA1 region of the dHPC. Prior to surgery, rats were anesthetized using isoflurane (Benson Medical Industries, Markham, Ontario). Following that, the head was shaved and a vertical incision (3-4 cm) was made to expose the skull. The skin was retracted and Bregma visualized using Q-tips covered with hydrogen peroxide. Following the measurement of Bregma, four screws were drilled in opposing ends of the skull and a guide cannula was dropped at the designated coordinates for area CA1 relative to Bregma (Paxinos and Waston, 2007); anterior-posterior -3.8mm, medial/lateral: ±2.5mm, and dorsal/ventral: -2.5mm. The cannula was then stabilized using a combination of dental cement and jet liquid. Dummy cannulas (0.36 mm) were inserted into the guide cannulas before the rat was placed in a recovery cage under heat for a period of 30 min. Once the rat was awake and bright, alert, and responsive, he was moved to his home cage to recover for a minimum of 7 d.

**Microinfusion Procedure**

Micro-infusions took place in a room separate from where behavioural testing was performed. The dummy cannulas were removed while the rat was gently restrained and 28 gauge infusion cannulas were inserted into the guide cannula. Bilateral infusions were conducted using two Hamilton syringes connected to a Harvard apparatus precision pump (Hillston, Massachusetts). The infusion process lasted for 2 min. Following this, the infusion cannulas were removed, and the dummy cannulas were reinserted. All animals were habituated to the infusion process for 2 days prior to the training procedure commencing; during these sessions, dummy cannulas were removed and the infusion cannulas inserted, but no fluid was infused.

### **Drug Administration: Intra-cranial**

Pirenzepine (Sigma-Aldrich, Oakville, Ontario), a selective M1 mAChR antagonist was given in experiment 5. It was administered at a dose of 20 μg/μL 30 minutes prior to reactivation, which has previously been shown to block destabilization of object location memories in dHPC (Huff et al., 2022). Physiological saline (0.9%) was used as a control (VEH) in each experiment.

**Fear Conditioning Apparatus**

Four fear conditioning boxes (30cm x 24cm x 24cm), each housed within separate sound-attenuating chambers (Med Associates Inc., Fairfax, VT), were used for all experiments. Near infrared imaging was accomplished via a camera mounted inside the door of each chamber. The camera captured movement at 30 frames per second and automatically scored freezing through Video Freeze software (Med Associates Inc., Fairfax, VT). Throughout the experiments, the house light was always on in the boxes. Between rats, boxes were cleaned with 5 % hydrogen peroxide (H2O2).

**Data Analysis**

All freezing behavior was collected using Med Associates Inc’s stock software, Video Freeze. Freezing was defined as no movement except that required for breathing (Maren, 2001). Freezing was automatically scored by the software in real time and verified offline by the researcher for discrepancies. Raw data files were analyzed using Video Freeze’s internal component analysis function, which bases each analysis on the session’s duration and parameters. Freezing behavior was denoted as percent freezing out of 100% based on the timing of each session. Freezing behaviour in alternate contexts was recorded using a video camera situated above each of the alternate contexts and was analyzed using ezTrack, a freezing analysis software (Pennington et al., 2019). One-way analysis of variance (ANOVA) was used to determine main group effects for experiments 1-4. For experiment 5, a 2x2 ANOVA was used. Tukey post-hoc tests were conducted for multiple comparisons. In addition, independent samples t-tests were used for experiments with two groups. All statistical analyses were conducted with IBM SPSS (Version 28).
